# Improved prediction of Bovine Leucocyte Antigens (BoLA) presented ligands by use of MS eluted ligands and in-vitro binding data; impact for the identification T cell epitopes

**DOI:** 10.1101/195016

**Authors:** Morten Nielsen, Tim Connelley, Nicola Ternette

## Abstract

Peptide binding to MHC class I molecules is the single most selective step in antigen presentation and the strongest single correlate to peptide cellular immunogenicity. The cost of experimentally characterizing the rules of peptide presentation for a given MHC-I molecule is extensive, and predictors of peptide-MHC interactions constitute an attractive alternative.

Recently, an increasing amount of MHC presented peptides identified by mass spectrometry (MS ligands) has been published. Handling and interpretation of MS ligand data is in general challenging due to the poly-specificity nature of the data. We here outline a general pipeline for dealing with this challenge, and accurately annotate ligands to the relevant MHC-I molecule they were eluted from by use of GibbsClustering and binding motif information inferred from *in-silico* models. We illustrate the approach here in the context of MHCI molecules (BoLA) of cattle. Next, we demonstrate how such annotated BoLA MS ligand data can readily be integrated with *in-vitro* binding affinity data in a prediction model with very high and unprecedented performance for identification of BoLA-I restricted T cell epitopes.

The approach has here been applied to the BoLA-I system, but the pipeline is readily applicable to MHC systems in other species.

## Introduction

Binding to MHC class I molecules (MHC-I) is a prerequisite for antigen presentation and induction of cytotoxic T cell responses ^1^. MHC-I molecules are highly specific, binding only a very small part of the possible peptide space. This high specificity combined with its essential role of antigen presentation has placed MHC in the focal point of research related to rational T cell epitope discovery and vaccine design.

Traditionally, the specificity of MHC molecules has been characterized using *in-vitro* binding assays ^2^. Using such assays, binding affinity values for large sets of single peptides have allowed very accurate experimental characterization of the binding motifs for a large panel of MHC-I molecules including the most prevalent human and several non-human primate MHC-I molecules. Recent studies have further extended the approach to livestock species including pig (SLA) ^3^ and cattle (BoLA) ^4^. However, the cost of applying an *in-vitro* approach to the characterization of MHC-I molecules is substantial and using it to characterise all MHC molecules within a given species remains unfeasible.

Given this, large efforts have been dedicated to the development of accurate *in-silico* models capable of characterizing the specificity of MHC-I molecules that not only allow the prediction of peptide binding to MHC-I molecules outside the very small set of peptides with measured binding affinity (i.e. extrapolating the peptide-space) but also make predictions for MHC-I molecules with limited or even no experimental binding data (i.e. extrapolating the MHC-I space). One method capable of both these types of extrapolations is NetMHCpan ^5,6,7^. This method is pan-specific in the sense that it allows prediction of peptide binding to any MHC-I molecule with known protein sequence, and the method has in several benchmark studies been shown to be ‘state-of-the-art’ ^8^.

One inherent problem with most MHC-I binding prediction methods available, including NetMHCpan, is that they reflect the nature of the data used in the training of the underlying models. Since most prediction methods are trained on in-vitro binding data, the predictive power of the models is restricted by any bias present in the in-vitro binding data. It is clear that several biases are present in the currently available in-vitro binding data, compromising their relevance as a descriptor of the biological event of antigen presentation: Binding data does not address the fact that antigen presentation is a complex integrative physiological process that combines antigen processing and transport as well as binding affinity and binding stability of the peptide to the MHC-I binding groove. Additionally, in vitro data fails to reflect any peptide length preference of different MHC-I alleles.

Recent advances in liquid chromatography tandem mass spectrometry (LC-MS2) have allowed this technique to become a powerful alternative to *in-vitro* binding assays for the identification of MHC ligands, T cell epitopes ^9,10^ and characterization of MHC binding specificities ^11^. The use of LC-MS^2^ to identify MHC ligands has several clear advantages over *in-vitro* binding assays. First and foremost the data obtained suffers to a much lesser degree from the biases described above for the *in-vitro* generated binding data. However, a limiting factor of the LC-MS^2^ approach for identification of MHC ligands is the sensitivity with which peptides can be reliably identified. It is inherent to the technology that the most abundant peptides in the sample are prioritised for identification using standard LC-MS^2^ acquisition methods. This is hallmarked by the fact that only a minor proportion of known T cell epitopes are identified as MHC ligands in standard LC-MS^2^ assays. Hence, in the foreseeable future, LC-MS^2^ will, in line with *in-vitro* assays, serve as a guide and not a solution for rational identification of T cell epitopes.

Recent studies have suggested that training prediction engines on LC-MS^2^-determined MHC ligand data (further referred to as MS ligand data) rather than binding affinity data improves the ability to accurately identify MHC ligands ^11,12,13^. This observation strengthens the assumption that MS ligand data may provide a better representation of the presented peptide antigen pool compared to *in-vitro* binding affinity data.

However, since the number of different MHC-I molecules characterized by MS ligands compared to *in-vitro* binding data remains small, we have in a recent publication outlined an approach that permits the inclusion of both MS ligand data and binding affinity data into a framework for learning MHC-I peptide interactions ^8^. In this earlier work, we demonstrated how this approach increased predictive performance compared to the previous ‘state-of-the-art’ techniques with regards to identification of naturally processed ligands, cancer neo-antigens, and T cell epitopes.

Here, we extend the approach to livestock. We outline a general pipeline for analysis and interpretation of MS ligand data. The pipeline consists of three steps; i) clustering of MS ligand data into MHC specificity groups, ii) association of specificity groups to specific MHC molecules, and iii) integration of MS ligand data into a framework for prediction of MHC-restricted T cell epitopes. We describe this pipeline in the context of bovine leukocyte antigen class I (BoLA-I) ligand data, and demonstrate how the approach can readily be applied to construct a predictive model with very high and unprecedented performance for identification of BoLA-I restricted T cell epitopes.

## Material and methods

### BoLA-I-associated peptide purification

Cells were washed and then lysed using 10 ml lysis buffer (1% Igepal 630, 300 mM NaCl, 100 mM Tris pH 8.0 and protese inhibitors) per 10^9^ cells. Lysates were cleared by centrifugation at 500 *g* for 10 min followed by 15,000 *g* for 60 min. BoLA complexes were captured using a pan anti-BoLA class I antibody IL-88 that was covalently conjugated to protein A sepharose immunoresin (GE healthcare) at a concentration of 5mg/ml. Bound complexes were washed sequentially using buffers of 50 mM Tris buffer, pH 8.0 containing 150 mM NaCl, then 400 mM NaCl, and finally 0mM NaCl. BoLA-associated peptides were eluted using 5 ml 10% acetic acid and dried.

### High performance liquid chromatography (HPLC) fractionation

Affinity column-eluted material was re-suspended in 120 μl loading buffer (0.1% formic acid, 1% acetonitrile in water). Samples were loaded onto on a 4.6 x 50 mm ProSwiftTM RP-1S column (Thermo Scientific) and eluted using a 500 μl/min flow rate over 10 min from 2-35 % buffer B (0.1% formic acid in acetonitrile) in buffer A (0.1% formic acid in water) using an Ultimate 3000 HPLC system (Thermo Scientific). Sample fractions were collected from 2-15 min. Protein detection was performed at 280 nm. Fractions up to 12 min that did not contain ß_2_-microglobulin were combined, dried and further analysed by LC-MS^2^.

### LC-MS^2^ analysis

Samples were suspended in 20 μl loading buffer and analysed on an Ultimate 3000 nano UPLC system online coupled to a QExactive-HF mass spectrometer (Thermo Scientific). Peptides were separated on a 75 μm x 50 cm PepMap C18 column using a 2h linear gradient from 5% buffer A to 35% buffer B at a flow rate of 250 nl/min (approx. 600 bar). Peptides were introduced into the mass spectrometer using a nano Easy Spray source (Thermo Scientific). Subsequent isolation and higher-energy C-trap dissociation (HCD) was induced on the 20 most abundant ions per full MS scan with an accumulation time of 128 ms and an isolation width of 1.0 Da. All fragmented precursor ions were actively excluded from repeated selection for 8 s.

### MS data analysis

Sequence interpretations of MS^2^ spectra were performed using a database containing all bovine UniProt entries combined with the all annotated *Theileria parva* proteins (35,992 entries total; bovine SwissProt entries: 5,995, bovine Trembl entries: 25,911, *Theileria parva* entries: 4,084). Spectral interpretation was performed using PEAKS 7.5 (Bioinformatics Solutions Inc.).

### NetMHCpan retraining and data preparation

MHC binding affinity data were obtained from the IEDB ^14^ (http://tools.immuneepitope.org/main/datasets; dataset used for retraining the IEDB class I binding prediction tools). This dataset consists of 186,684 peptide-MHC binding affinity measurements covering 172 MHC molecules from human, mouse, primates, cattle, and swine. IEDB eluted ligands were also obtained from the IEDB applying the filtering procedure described in ^8^. This data set contains 85,217 entries in total restricted by 55 unique MHC molecules. Random artificial negatives were added for each MHC molecule covered by eluted ligand data by sampling randomly 10*N peptides of each length 8-15 amino acids from the antigen source protein sequences, where N is the number of 9mer ligands for the given MHC molecule. A similar procedure was applied for the BoLA-I restricted eluted ligands. Here, however the artificial negatives were obtained from a set of random natural proteins.

## Results

### The raw MS ligand data

We first analysed the accuracy and consistency of the raw MS BoLA-I ligand data generated. MS ligand data were obtained from 5 BoLA-I homozygous cell lines describing 3 BoLA haplotypes; A10, A14, and A18. More than 94% of the peptides identified had lengths of between 8 and 14 amino acids. Focusing on this peptide subset, the number of ligands obtained in each experiment varied between 6,615 and 7,755 (Fig. 1A). The overlap in ligands identified between cell-lines expressing the same BoLA-I haplotype was large, with more than 50% of the unique peptides for each haplotype data set found in both samples (reflected in the total number of ligands for each haplotype being smaller than the sum of the counts from each cell line) (Fig. 1B).

**Figure 1.**
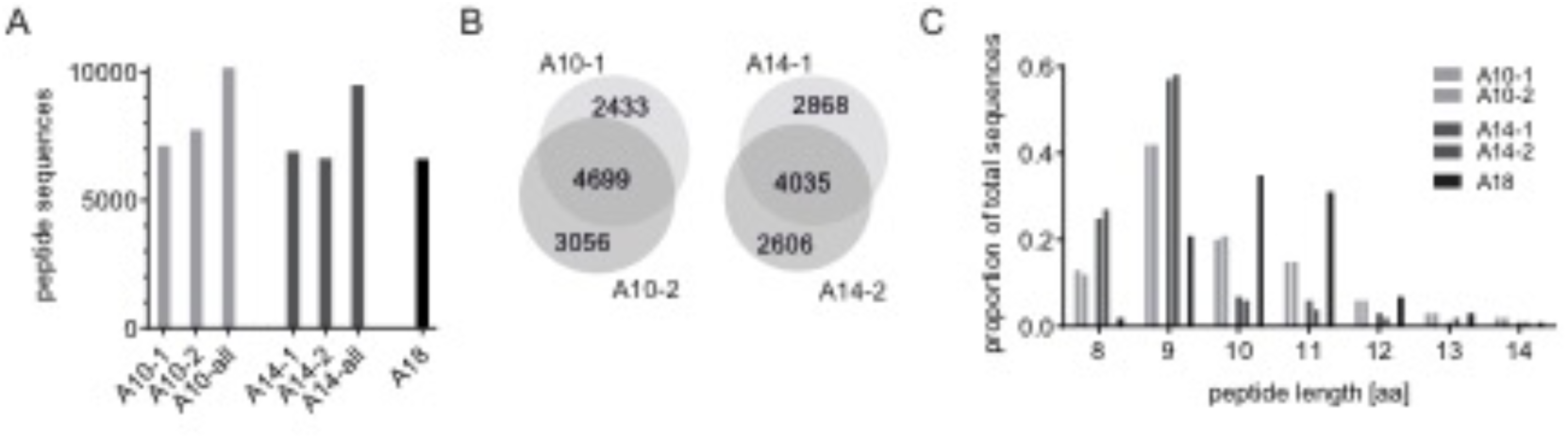
(A) Number of peptides obtained from each cell line, and for the combined set (all) of peptides for each of the BoLA-A10, -A14 and –A18 haplotypes. (B) Overlap of peptide sequences for A10 and A14 samples. (C) Distribution of ligand lengths within the different data sets.

The length distribution of the peptides was highly consistent between data sets from each haplotype, but varied substantially between haplotypes (Fig. 1C). The extreme examples being the data from cell lines expressing the A14 haplotype, with a relatively high preference for 8mers, and the cell lines expressing the A18 haplotype, with a preference for 10 and 11mer peptides.

### Identification of MHC-I allele specificity groups

One first challenge faced when interpreting and analysing MS ligand data obtained from a given cell line that expresses more than a single MHC allele, is assigning the peptides to the relevant MHC-I molecule they were eluted from. We have earlier demonstrated that this challenge can be readily and accurately solved using the GibbsCluster method ^15^. In short, this method takes the complete list of eluted ligands from a given experiment as input, and seeks to cluster the ligands into groups so that similarity within each group is high and the similarity between groups is low. The algorithm includes an option to place ligands into a trash cluster if they demonstrate poor similarity to all defined clusters. This trash cluster option has proven very powerful for removal of false positive ligand data. The recent update to the tool allows the clustering to be performed on peptides of variable length ^16^. The outcome of the algorithm is a solution defined by an optimal number of clusters, with each ligand associated to one such cluster (or the trash cluster).

We hence applied the GibbsCluster method (version 2.0) to deconvolute the BoLA-I restriction of the ligands in the data sets corresponding to the three haplotypes. The method was run with default options for MHC class I ligand clustering except for the number of seeds, which was set to 10 to allow for improved sampling (Fig. 2).

**Figure 2.**
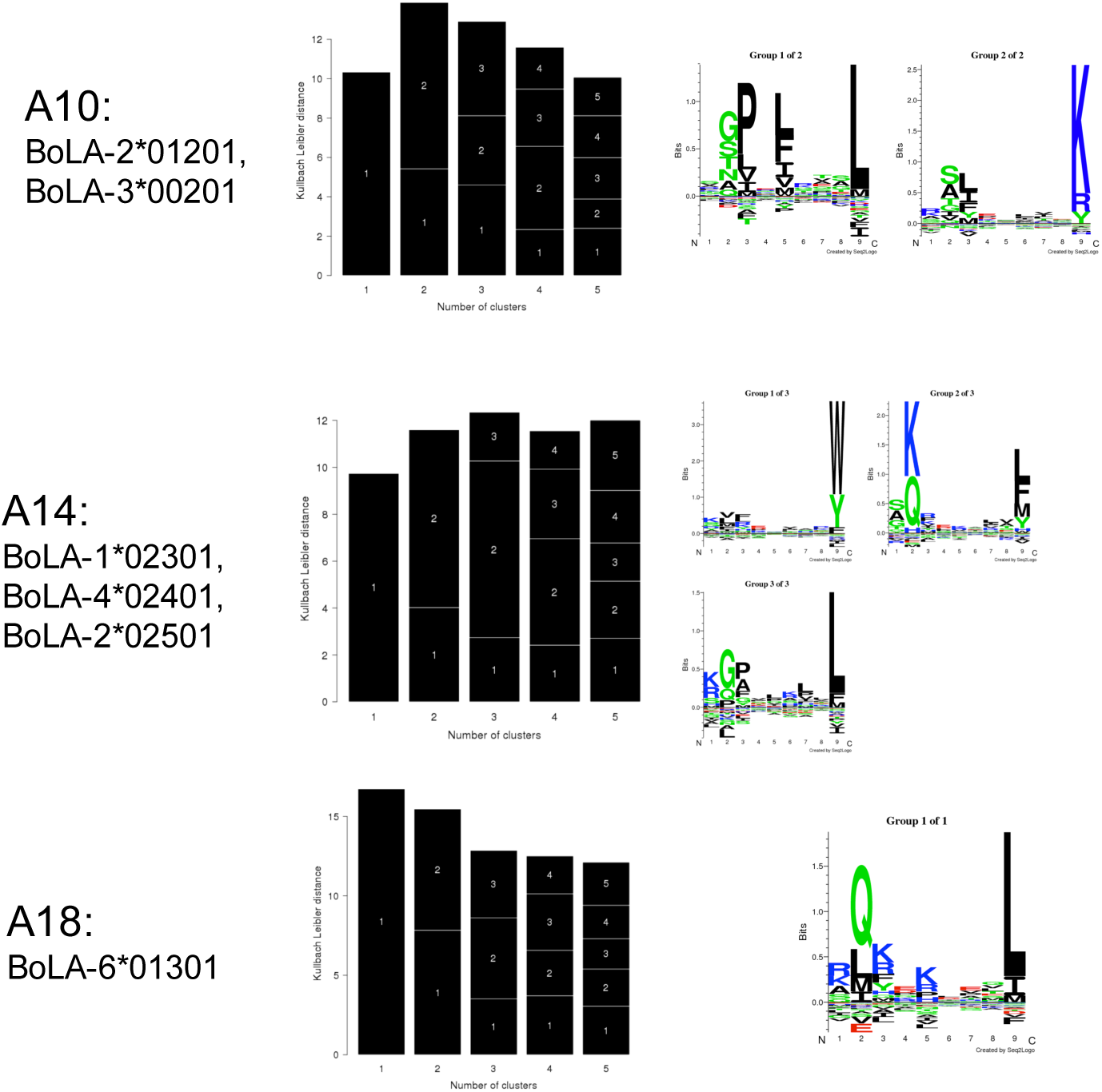
GibbsCluster analysis of the three combined data sets. Each row displays the results from one haplotype data set. Left panels give the barplot of the Kullback-Leibler Distance (KLD) as a function of the number of clusters. The relative size of each black block within a bar is proportional to the size of each of the clusters. The right panels give the sequence motifs derived from the best solution (i.e. the solution with highest KLD) displayed in the form of sequence logos generated with Seq2Logo ^17^.

The GibbsCluster method selects the solution with the highest KLD value (the central column of Fig. 2) and in all three cases, we find a perfect correspondence between the number of known functional BoLA-I alleles expressed by each cell line and the number of clusters identified by the GibbsCluster method. The number of peptides assigned to the trash cluster was in the three cases; 3.4% (BoLA A10: 347 out of 10,188), 2.1% (BoLA A14: 201 out of 9,509), and 2.5% (BoLA A18: 164 out of 6,615). These low proportions confirm the very high purity of the eluted ligand data sets.

### Annotation of BoLA-I restrictions to specificity groups

The next challenge of analysing the MS ligand data set is the association of each identified ligand clusters to a BoLA-I molecule expressed in the given cell line. Here, we used the motifs predicted by NetMHCpan (version 3-0) ^7^ as a qualitative guide to make this association. This approach allowed in all cases a clear and unambiguous association of a single BoLA-I restriction to each cluster (Fig. 3).

**Figure 3.**
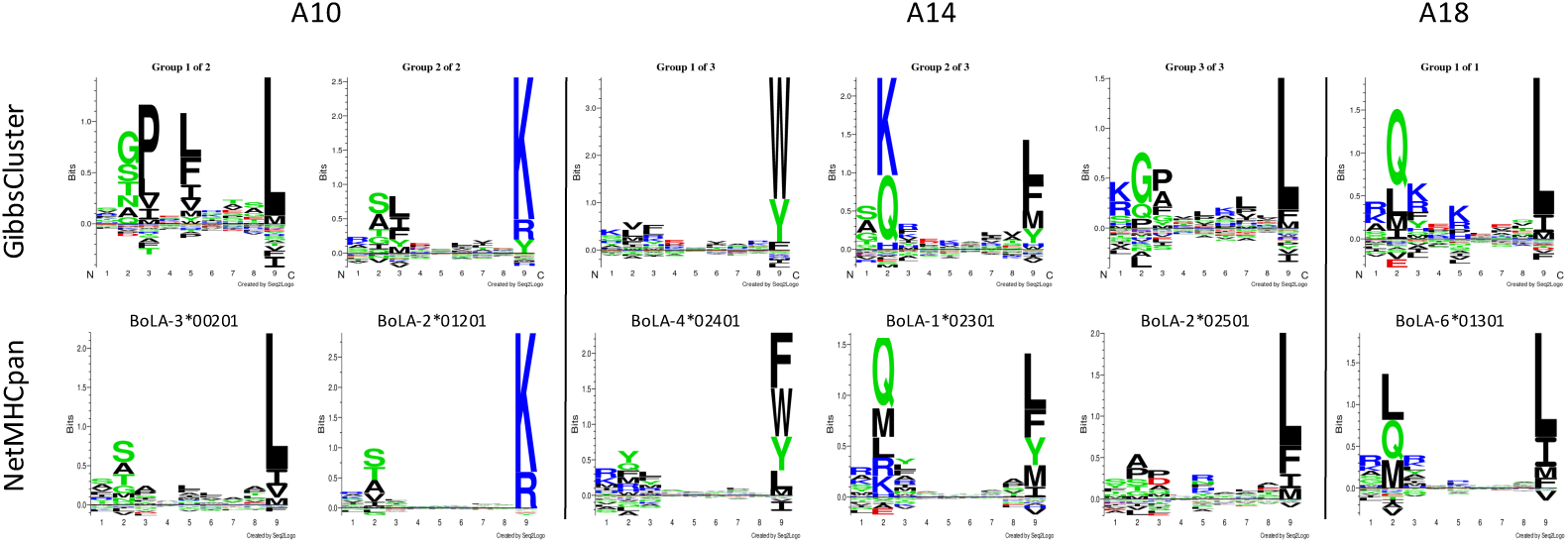
Mapping of GibbsClustered peptides to BoLA-I molecule specificities. Each haplotype is shown separated by the vertical lines as indicated. In each column the binding motif logos for each of the optimal GibbsCluster solutions (upper row) together with the best matched NetMHCpan predicted binding motif for the BoLA-I molecules (lower row) expressed by the relevant haplotype are shown (as determined by visual comparison).

Based on this mapping, we unambiguously assigned each cluster to one BoLA-I molecule as shown in Table 1.

**Table 1.** Association of GibbsCluster clusters to BoLA-I restrictions.

| Cell line | Group | BoLA-I |
| --- | --- | --- |
| A10 | G1 | BoLA-3*00201 |
|  | G2 | BoLA-2*01201 |
| A14 | G1 | BoLA-4*02401 |
|  | G2 | BoLA-1*02301 |
|  | G3 | BoLA-2*02501 |
| A18 | G1 | BoLA-6*01301 |

Given this mapping of ligands to individual BoLA molecules we were able to conduct an allele-specific analysis of the length distribution of MHC-I ligands. As expected most of the BoLA molecules had a length preference for 9mer peptides (Fig. 4). However, clear differences in the ligand length distribution between the different BoLA-I molecules was evident. The most extreme cases were BoLA-1*02301 (A14), with a high preference for binding 8mer peptides (>30%), and BoLA-6*01301 (A18) with an increased preference for binding 10 and 11mer peptides (>65%).

**Figure 4.**
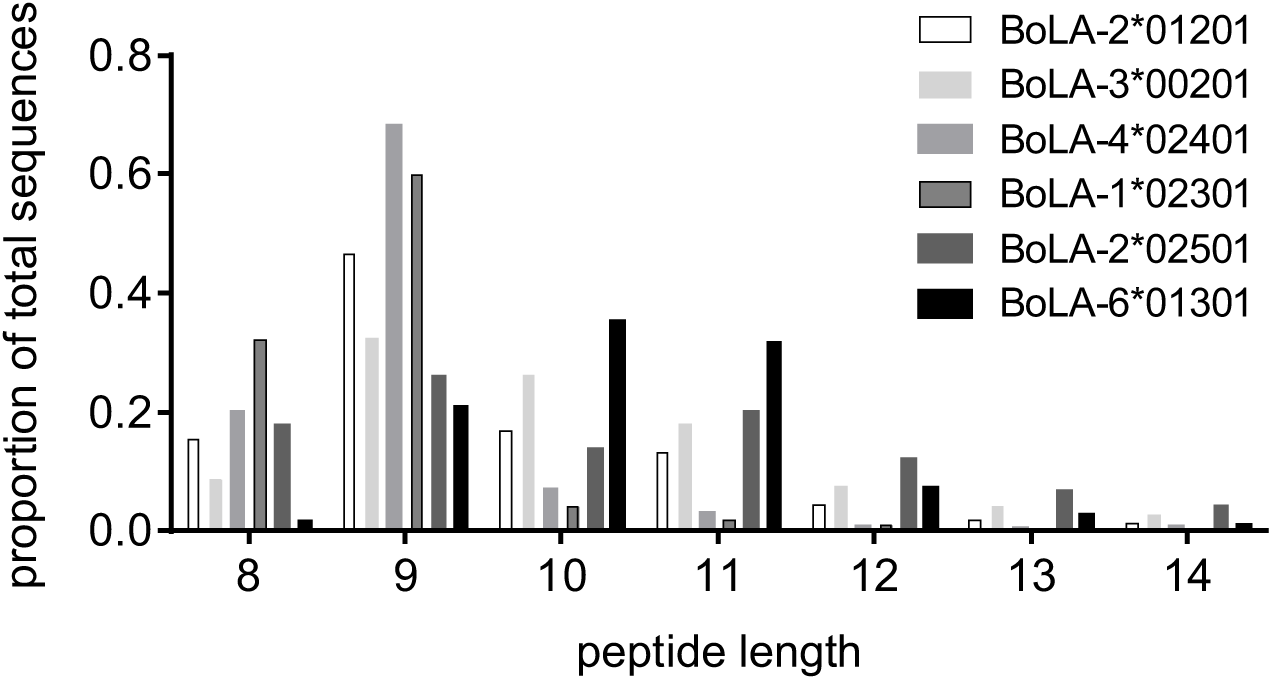
Length distribution of ligands restricted to each BoLA molecule.

### Construction of a prediction model

Having mapped the likely BoLA-I restriction of the individual ligands, we sought to use this information to (re)train the NetMHCpan prediction method combining the BoLA MS eluted ligand (EL) data with BoLA binding affinity (BA) data contained within the IEDB. The retraining of the NetMHCpan prediction method was performed as described earlier ^8^. In short, the method was trained on the two data types (BA and EL) in a conventional three-layer feed forward artificial neural network. The weights between the input layers and the hidden layer are shared between the two input types, and the output layer has two output neurons; one for each input type. During training, examples of the two data types (EL and BA) are shown at random to the network, and weights are adjusted using stochastic gradient descent along the gradient of the output neuron corresponding to the input type. Variations in peptide length are handled allowing insertions and deletions as described in ^18^.

The IEDB binding affinity peptide data set for 7 BoLA molecules (listed in table 2) contains exclusively 9mer peptide data. By integrating the MS ligand, we would hence expect that the updated NetMHCpan method would achieve an improved predictive performance for the BoLA system due to i) the method being informed of the differences in length preference of the different BoLA molecules as illustrated in Fig. 4, and ii) the inclusion of peptide data for additional BoLA molecules expanding the knowledge of BoLA binding specificities. Examples of i) are shown in Fig. 5. Here, the top 1% percent strongest predicted binders from a set of 700,000 random natural 8-14mer peptides (100,000 of each length) predicted using *NetMHCpan-2.8, NetMHCpan-3.0*, and the two individual output values of the method trained on the combined BoLA-eluted MS ligand data (EL) and binding affinity data (BA) (method 4.0) are shown in comparison to the measured length distribution of the MS ligand data.

**Table 2.** Comparison of the predictive performance of NetMHCpan-4.0_BA (the binding affinity prediction score of the NetMHCpan-4.0 method trained on both eluted ligand and peptide binding affinity data) and NetMHCpan-3.0 models on quantitative binding affinity data from the IEDB affinity data set. Names in parentesis in the first column refer to the historial names for the dfferent alleles. Performance was estimated in terms of Pearson’s correlation coefficient (PCC) and AUC (area under the receiver operator curve). Both these performance measures take a value of 1 for the perfect, and values of 0.0 (PCC)/0.5 (AUC) for a random prediction.

| BoLA-I | #peps | #bind | NetMHCpan-4.0_BA |  | NetMHCpan-3.0 |  |
| --- | --- | --- | --- | --- | --- | --- |
|  |  |  | PCC | AUC | PCC | AUC |
| BoLA-3*00101 (BoLA-AW10) | 166 | 8 | 0.497 | 0.816 | 0.381 | 0.792 |
| BoLA-1*02301 (BoLA-D18.4) | 258 | 182 | 0.648 | 0.832 | 0.551 | 0.747 |
| BoLA-6*01301 (BoLA-HD6) | 268 | 219 | 0.622 | 0.815 | 0.482 | 0.728 |
| BoLA-3*00201 (BoLA-JSP.1) | 158 | 32 | 0.464 | 0.703 | 0.277 | 0.622 |
| BoLA-T2c | 90 | 84 | 0.485 | 0.833 | 0.455 | 0.813 |
| BoLA-2*01201 (BoLA-T2a) | 167 | 47 | 0.691 | 0.852 | 0.635 | 0.812 |
| BoLA-6*04101 (BoLA-T2b) | 157 | 38 | 0.631 | 0.835 | 0.566 | 0.816 |
| Ave |  |  | 0.577 | 0.812 | 0.478 | 0.761 |

**Figure 5.**
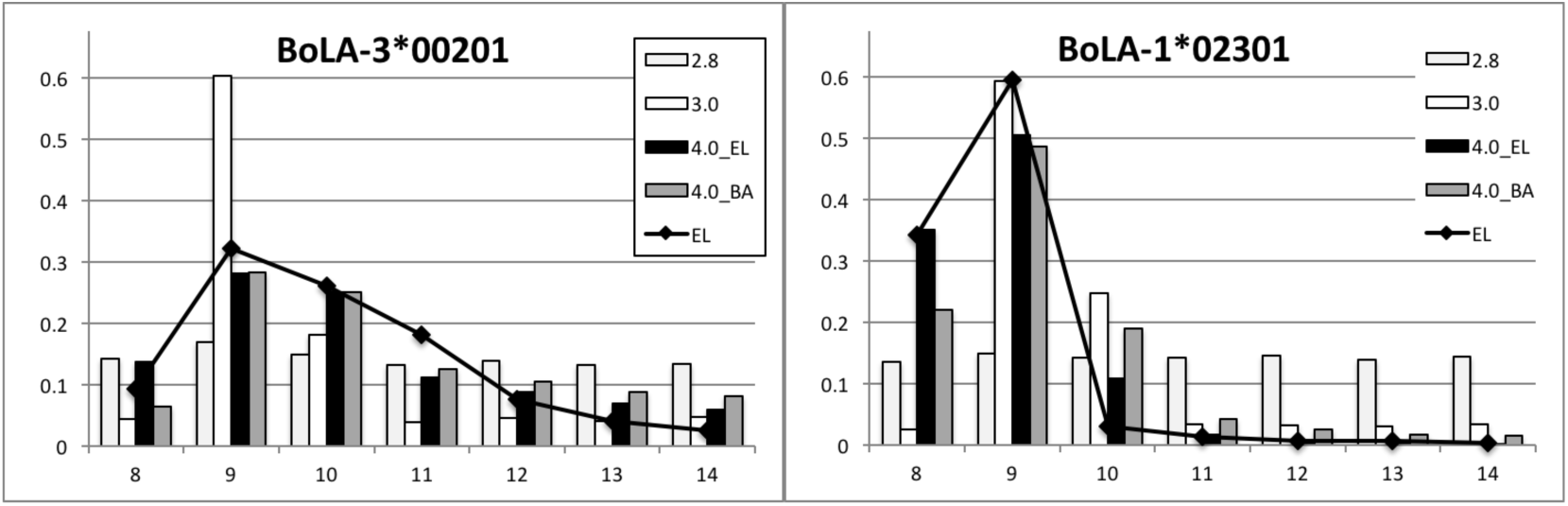
Predicted length preference for the BoLA-3*00201 (left) and BoLA-1*02301 (right) molecules. The solid line show the length distribution for the MS eluted ligands in both panels and bars shown the length distribution predicted by NetMHCpan-2.8 (2.8 - light grey), NetMHCpan-3.0 (3.0 - white), the eluted ligand output value of NetMHCpan-4.0 (4.0_EL - black), and the binding affinity output value of NetMHCpan-4.0 (4.0_BA - grey).

The figure shows that the MS eluted ligand likelihood score (4.0 EL) and to a lesser degree the binding affinity score (4.0 BA) of the NetMHCpan-4.0 model trained on the combined BoLA-eluted MS ligand and IEDB MHC binding affinity data accurately captures the length preference of the two BoLA molecules as described from the MHC-elution MS ligand data. The figure also confirms earlier observation that NetMHCpan-2.8 has no power to predict peptide length preferences of different MHC molecules, and that *NetMHCpan-3.0* in most cases (included the two shown here) predicts a strong preference for 9mers followed by 10mers but has very limited predictive potential for peptides of other lengths ^7^.

Turning next to the evaluation of how the inclusion of the MS ligand data impacts the performance when it comes to predicting peptide-BoLA interactions, we show in Table 2 the performance of models trained with and without MS ligand data (NetMHCpan-3.0 and NetMHCpan-4.0, respectively) when evaluated on the set of peptides with measured binding affinity data from the IEDB (performance is evaluated using 5-fold cross validation). From this evaluation, it is apparent that adding MS ligand data improves the predictive performance of the model with a consistent increase in the predictive power as measured in terms of both the Pearson’s correlation coefficient (PCC) and area under the receiver operator curve (AUC) across all 6 BoLA molecules.

### Identification of BoLA-I restricted T cell epitopes

Finally, we evaluated the combined impact of the above two effects (improved prediction of the preferred peptide length, and expanded knowledge of BoLA binding specificities) for the capacity to enhance prediction of BoLA-I restricted T cell epitopes. We performed this evaluation on a set of previously described epitopes with known BoLA-I restrictions ^19,20^. Here, we predict binding for all overlapping 8-11mer peptides from the source protein of the epitopes to the known BoLA-I restriction molecule using the NetMHCpan-3.0 and NetMHCpan-4.0_EL (the eluted ligand prediction score of the NetMHCpan-4.0 method trained on both eluted ligand and peptide binding affinity data) prediction methods, and report the performance of each method as a Frank score (i.e. the fraction of peptides with predicted binding values greater than the epitope). Using this measure, a value of 0 corresponds to a perfect prediction (the known epitope is identify with the highest predicted binding value among all peptides found within the source protein), and a value of 0.5 to random prediction. The result of this evaluation is shown in Table 3

**Table 3.**
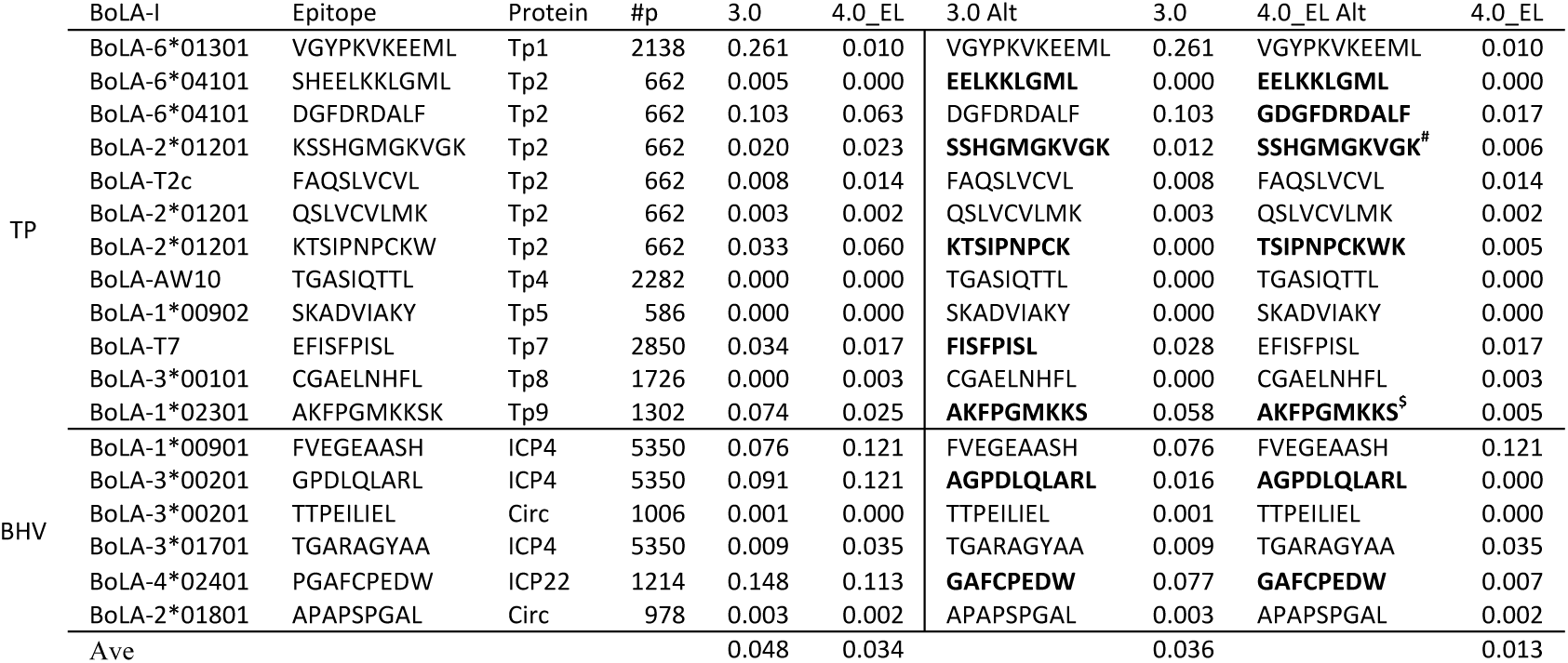
Predictive performance of the NetMHCpan-4.0 eluted ligand likelihood prediction model (4.0_EL) compared to NetMHCpan-3.0 (3.0) on a data set of known BoLA-I restricted T cell epitopes from Theileria parva (TP) and Bovine herpes virus (BHV). The part of the table to the left of the vertical line gives the performance of the two methods on the original epitope data. The part of the table to the right of the vertical line gives the results allowing each prediction method to suggest alternative epitopes overlapping with the known epitopes (either contained within known epitopes or with single amino acid extensions). In bold is high lighted the case where the two methods suggest alternative optimal epitopes. #Minimal epitope defined in 21, $Minimal epitope (N. MacHugh personal communication).

Also in this data, the improvement in predictive performance when integrating the BoLA-MS eluted ligand data in the prediction methods is apparent. When permitting the prediction methods to suggest alternative epitopes the average scores for NetMHCpan3.0 and 4.0_EL are 0.036 and 0.013 respectively. In the majority of cases (16/18), the NetMHCpan-4.0 method identifies the epitopes (or suggests an alternative variant) within the top 2% of peptides contained within the source protein of the epitope (i.e. has a Frank value less than 0.02). Notably, the worst prediction made by NetMHCpan4.0_EL is for the FVEGEAASH epitope presented by BoLA-1*00901; there is no binding or BoLA-eluted ligand data are available characterizing the specificity and ligand length preference of this BoLA-I molecule. The closest neighbour, defined in terms of the sequence similarity of the pseudo sequence of BoLA-1*00901 to the MHC molecules in the training data, is BoLA-1*02301 with a neighbour distance of 0.14. This value is larger than the 0.1 distance value that as rule of thumb is defined as the threshold for when NetMHCpan predictions are reliable ^22^. It is hence not unexpected to observe low predictive performance for this molecule and its exclusion nearly halves the average score for the NetMHCpan4.0_EL predictions to 0.007. In conclusion, in practical terms and in the context of workload reduction, this result means that more than 99% of the peptides contained within the antigen protein sequences could be discarded by use of peptide binding prioritizations prior to experimental validation.

## Discussion and Conclusions

In this work, we have outlined a simple yet highly powerful pipeline for the analysis and interpretation of LC-MS^2^-defined MHC-eluted ligand data. We applied the pipeline to analyse eluted ligand data obtained from 5 cell lines covering 3 BoLA-I haplotypes each characterized to express between one and three distinct BoLA-I molecules.

We demonstrate how the pipeline can effectively deal with several of the important challenges when interpreting MS ligand data, including identification of false positives, identification of the MHC binding motif and assignment to the regarding MHC restriction elements. We also demonstrate how this information can be integrated into prediction methods to improve their accuracy for rational epitope discovery.

Comparing the set of ligands identified from separate cell lines derived from two different animals expressing identical BoLA haplotypes revealed a high level of consistency with more than 50% of the ligands shared between both. Using the GibbsCluster method to group the ligands allowed for both the identification of false positives and also MHC binding specificity clusters. A very low number of false positive peptides were identified (less than 3.4% in all samples), confirming the high accuracy of the MS ligand data. For each of the analysed data sets, we found a perfect correspondence between the number of specificity clusters identified and the number of known functional MHC molecules included in the corresponding haplotype. Analysing each of the identified clusters revealed large differences not only in binding motif of associated ligands but also in the length preference of the ligands presented by each of different MHC molecule. This difference in ligand length preference between BoLA-I molecule has to the best of our knowledge not been characterized before.

A challenge related to the identification of MHC binding specificity clusters is the subsequent association of specificity clusters to restricting MHC molecules expressed by the given cell. Several approaches have been suggested to deal with these challenges including the use of MHC mono-allelic cell lines ^13^ and co-occurrence of MHC alleles across different samples ^12^. However, in the vast majority of cases, the association can readily be obtained by comparing the binding motif of each specificity cluster to the motif predicted by state-of-the-art MHC binding prediction methods such as NetMHCpan ^7^. It is clear that this approach is limited by the prediction accuracy of the NetMHCpan method, and will fail in situations where NetMHCpan has no predictive power for one or more of the MHC molecules in question. However, as shown here, applying this approach where some prior information about the BoLA-I specificities is available, allows clusters to be unambiguously assigned to BoLA-I molecules.

Having mapped each ligand to a specific BoLA-I molecule, we analysed how to best benefit from these data in terms of development of improved prediction methods for BoLA-I restricted T cell epitopes. We have earlier demonstrated that integrating binding affinity data covering a range of relevant BoLA-I into the NetMHCpan prediction tool led to an improved performance for the identification of known BoLA-I restricted T cell epitopes ^4^. Earlier work moreover suggested that the MHC ligand prediction method could benefit from the integration of MS elution data ^13,11,23^. To benefit from both of these observations, we here applied the recently proposed approach to train the MHC binding predictor on a combined data set of binding affinity and MS ligand data ^8^. In accordance with the work by Jurtz *el al.* ^8^, we find that this approach led to significantly improved performance both for prediction of peptide binding affinity, and T cell epitopes.

As mentioned earlier, the different BoLA-I molecules showed very different peptide length binding preference. As shown here, including the MS ligand data in the training, allows, in agreement with earlier work, the prediction method to learn these differences, and hence boost the predictive performance by placing lower binding values to peptides with atypical length according to the eluted ligand length profile.

Evaluating the prediction model trained on the combined binding affinity and eluted ligand data on a set of validated BoLA-I restricted epitopes, we found a consistent improvement in performance compared to current methods. In agreement with earlier studies, the vast majority of epitopes are identified within the top 2% of the predicted peptides contained within the antigen source protein ^7^. Only one epitope was very poorly predicted by the proposed model. This epitope is restricted to a BoLA-I molecule very distinct (in terms of the protein sequence) from the BoLA molecules included in the training of the NetMHCpan method, and this most likely accounted for the low prediction accuracy for this molecule ^22^. Further, data on the nature of the peptides that bind to this allele is likely to improve the predictive values of NetMHCpan. Also, the analysis suggests, in agreement with earlier studies, the presence of alternative epitopes overlapping with the known epitopes (either wholly contained within the peptide or accommodated by single amino acid extensions) strongly suggest that these need to be refined to map the minimal epitope ^20,21,24^.

In conclusion, the present study confirms the very high accuracy of ‘state-of-the art’ proteomic methods for high throughput and accurate identification of MHC-presented ligands, and demonstrates how the proposed pipeline combining GibbsClustering and advanced data mining techniques allows the intuitive interpretation of MS ligand data and also the integration of such data for improved prediction of MHC peptide binding and T cell epitopes.

Here, the approach has been applied to the BoLA-I system, but the pipeline is readily applicable to MHC systems in other species.

## Acknowledgement

Mass spectrometry analysis was performed in the Target Discovery Institute Mass Spectrometry Laboratory led by Benedikt M. Kessler. The work was supported by the Department for International Development of the United Kingdom and the Bill and Melinda Gates Foundation (OPP1078791) and the CGIAR Research Program on Livestock and Fish.

